# Transdiagnostic multimodal neuroimaging in psychosis: structural, resting-state, and task MRI correlates of cognitive control

**DOI:** 10.1101/284273

**Authors:** Dov B. Lerman-Sinkoff, Sridhar Kandala, Vince D. Calhoun, Deanna M. Barch, Daniel T. Mamah

## Abstract

**Background:** Psychotic disorders, including schizophrenia and bipolar disorder, are associated with impairments in regulation of goal-directed behavior, termed cognitive control. Cognitive control related neural alterations have been studied in psychosis. However, studies are typically unimodal and relationships across modalities of brain function and structure remain unclear. Thus, we performed transdiagnostic multimodal analyses to examine cognitive control related neural variation in psychosis.

**Methods:** Structural, resting, and working memory task imaging and behavioral data for 31 controls, 27 bipolar, and 23 schizophrenia patients were collected and processed identically to the Human Connectome Project (HCP), enabling identification of relationships with prior multimodal work. Two cognitive control related independent components (ICs) derived from the HCP using multiset canonical correlation analysis + joint independent component analysis (mCCA+jICA) were used to predict performance in psychosis. *de novo* mCCA+jICA was performed, and resultant IC weights were correlated with cognitive control.

**Results:** *A priori* ICs significantly predicted cognitive control in psychosis (3/5 modalities significant). *De novo* mCCA+jICA identified an IC correlated with cognitive control that also discriminated groups. Structural contributions included insular, somatomotor, cingulate, and visual regions; task contributions included precentral, posterior parietal, cingulate, and visual regions; and resting-state contributions highlighted canonical network organization. Follow-up analyses suggested *de novo* correlations with cognitive control were primarily influenced by schizophrenia patients.

**Conclusions:** *A priori* components partially predicted performance in transdiagnostic psychosis and *de novo* analyses identified novel contributions in somatomotor and visual regions in schizophrenia. Together, results suggest joint contributions across modalities related to cognitive control across the healthy-to-psychosis spectrum.

## Introduction

Psychosis has been classically conceptualized as a hallmark of schizophrenia (SZ) (1), nonetheless it is present in a number of other disorders including schizoaffective and bipolar (BP) disorders. Importantly, alterations in cognition (2–4), including cognitive control (5, 6), are a key feature of psychosis. Furthermore, cognitive control alterations are observed transdiagnostically (7), including in individuals with SZ and BP (8), and other disorders (9–11). In the present work, we expand beyond prior work by using multimodal image analysis to examine transdiagnostic patterns of neural variation in structural, resting-state, and task functional imaging related to cognitive control.

Cognitive control refers to the set of cognitive functions that enable and support goal-directed behavior and regulation of one’s thoughts and actions (12), including the ability to maintain information over time (e.g., working memory), protect against distraction, and combine novel inputs (12, 13). Within the psychosis spectrum, SZ have poorer cognitive control performance than healthy controls (HC), and BP often have intermediary performance between SZ and HC (7, 14, 15). This graded performance further supports conceptualizations of psychosis as a dimensional transdiagnostic construct.

The imaging literature has identified transdiagnostic neural alterations in psychosis related to cognitive control. For example, with structural imaging, working memory performance was inversely correlated with cortical thickness in the right rostral anterior cingulate and positively correlated with surface area in the left rostral anterior cingulate and right rostral middle frontal region in a cohort of individuals with either SZ or BP (16). A recent meta-analysis of structural alterations in a broad mental illness sample identified significant psychosis-related gray-matter losses in medial prefrontal cortex, insula, thalamus, and amygdala and significant gray-matter increases in the striatum as compared to HC and non-psychotic mental illnesses (17). Finally, Shepherd et al. (18) examined a transdiagnostic psychosis cohort and dichotomized patients into high and low executive function groups based on two-back working memory performance. Compared to HC, the low executive function group exhibited decreased gray matter volume in bilateral superior and medial frontal gyri, and right inferior operculum and hippocampus.

Several approaches to resting-state imaging have identified transdiagnostic neural alterations (19, 20) related to general cognition (21–23), with fewer studies examining cognitive control (24). For example, trail making task performance in persons with SZ and BP was positively correlated with average connectivity strength between the whole brain and the left caudate, temporal occipital fusiform cortex / lingual gyrus, and left thalamus (21). Further, a network connectivity approach in another transdiagnostic cohort (25) identified a significant relationship between global efficiency of the cingulo-opercular network and cognitive and executive function, along with significant decreases in cingulo-opercular network efficiency in psychosis (25). Additionally, an independent component analysis (ICA) approach to resting-state data (23) identified significant decreases in connectivity between a fronto/occipital component and a combined anterior default mode and prefrontal component in individuals with psychosis (SZ and psychotic BP) and unaffected siblings. In the same study, decreased connectivity was also identified from a meso/paralimbic component to a sensory/motor component, but this was only observed in individuals with SZ and not psychotic BP. Furthermore, recent analyses from the same dataset identified nine abnormal networks, seven of which showed significant correlations with cognitive control including visual, working-memory, visuomotor integration, default mode, and frontoparietal control networks (24).

Several studies have examined transdiagnostic alterations in task imaging related to cognitive control. For example, Brandt et al. (26) used ICA decompositions of two-back working memory task and identified nine task-related components. Of those, three components with spatial distribution in frontal and parietal regions corresponding to working-memory networks showed significant graded hyperactivation pattern (SZ>BP>HC). In addition, Smucny et al. (7) used an a priori contrast approach to examine activation in the dorsolateral prefrontal cortex (DLPFC) and superior parietal cortex (SPC) during AX-CPT task in a transdiagnostic population. They identified significant graded task performance (HC>BP>SZ) as well as BOLD responses in DLPFC and partially significant responses in SPC.

As reviewed above, there have been a number of studies of neural alterations in groups that experience psychosis. However, much of this literature is unimodal (see (27) for a review of exceptions). Thus, it remains unclear how extant results relate across the modalities as well as across disease state. A multimodal analysis examining structural, resting state, and diffusion imaging and their relationships to a cognitive battery in SZ identified a relationship between overall cognitive impairment and variation in cortico-striato-thalamic circuitry (28). Additionally, an examination of three modalities derived from structural imaging in a transdiagnostic cohort (HC, BP, and SZ) identified putative associations between gray matter alterations and processing speed, working memory, and attention (29). However, these associations were not significant across multiple diagnostic categories within the same component and did not survive multiple comparison correction. Importantly, both studies did not focus specifically on cognitive control.

We recently used multiset canonical correlation analysis with joint independent component analysis (mCCA+jICA) to study multimodal neural correlates of cognitive control in the normative population (30). The mCCA+jICA framework was chosen as it flexibly identifies patterns across modalities and decomposes them into maximally spatially independent sources of variance (31). This has proven to be a powerful analytic framework and has been used to identify abnormalities in SZ (32) and also discriminate between HC, BP, and SZ (33). Using a community sample from the Human Connectome Project (HCP) (34), we identified relationships between two mCCA+jICA-derived multimodal patterns and individual differences in cognitive control performance (30). These two components included structural and functional contributions from the anterior insula, visual, and parietal regions as well canonical resting-state network structures. Importantly, the findings were replicable in an independent sample of participants from the HCP using both predictive and independent analyses.

The goal of the present study was to examine transdiagnostic multimodal neural alterations in psychosis, using the results of our previous work to guide analyses. To accomplish this, we recruited a transdiagnostic psychosis cohort comprised of HC, SZ, and BP. Participants were imaged using the same HCP-customized scanner and performed a subset of the same HCP tasks. We first assessed the replicability of our normative findings by predicting performance in the psychosis participants using the two components from our prior work that were significantly related to cognitive control. We then performed an independent mCCA+jICA decomposition of this new transdiagnostic dataset to identify novel patterns of alteration.

## Methods

### Participants

Participants were recruited from a broader study of neural alterations in psychosis and included healthy controls (HC), schizophrenia or schizoaffective disorder (SZ), and bipolar disorder (BP). Prior to image pre-processing, there were n=35 HC, n=36 BP, and n=31 SZ available. Of this, n=31 / 30* HC, n=27 BP, and n=23 SZ had the requisite data for inclusion in this study (*see *supplement*). Participants were recruited from clinical and community settings in Saint Louis. Participants had no substance use disorder in the prior six months, no clinically significant head trauma, and no neurological diseases. Patient participation criteria included: DSM-IV diagnosis of BP or SZ, age 18-30, and stable outpatient or partial hospital status. HC were recruited to have similar demographics (age, gender, parental level of education) as patients. HC participation criteria included: no history of DSM-IV psychotic disorder and no cognitive enhancing or psychotropic medication for prior three months. Study procedures were approved by the Washington University IRB and all participants gave written informed consent.

### Behavioral assessment

A composite measure of cognitive control was generated for each participant from their performance on three tasks (see *supplement* for details on each task): (1) In-scanner N-Back task; 2) Out of scanner letter N-Back task from the Penn Computerized Neuropsychological battery (PennCNP) (35); and, 3) Progressive matrices from the PennCNP. Performance on each task was individually Z-scored and averaged to generate the composite measure. This composite had a Cronbach’s alpha of 0.76, suggesting good internal consistency.

### Neuroimaging collection and pre-processing

Participants were scanned at Washington University in St. Louis using a customized Siemens “Connectome” scanner developed for the HCP (34). T1 and T2 were acquired at 0.7mm isotropic resolution and BOLD imaging were collected at 2mm isotropic resolution with 720ms TR. Data were pre-processed using the HCP pipelines (36) and further processed to generate the final imaging measures used in the present study as was done in (30). Three imaging modalities were used: 1) cortical thickness (sMRI); 2) resting state functional connectivity correlation matrices (rsfcMRI) generated using cortical (37), cerebellar (38), and subcortical parcels from Freesurfer; and, 3) the HCP N-back working memory task fMRI (tfMRI) – activation in the 2-back condition (see *supplement*).

### Relationship to a priori normative multimodal correlates of cognitive control

Our previous work using mCCA+jICA identified replicable multimodal patterns from two mCCA+jICA components that were significantly related to cognitive control in healthy participants in the HCP (30). We performed an analysis to determine whether patterns from these two components also predicted cognitive control in the present participants. Given that data were collected and processed identically, we directly applied the three relevant modalities from these two components from the prior study to the source data from the present cohort (referred to as HCP_C1_IC2 and HCP_C1_IC7). This generated a set of subject-specific weights for the present cohort corresponding to the extent to which the *a priori* components reflected of the present data (see *supplement*). These weights were then correlated with the composite cognitive control measure.

### mCCA+jICA multimodal imaging analysis

We also identified mCCA+jICA components *de novo* in the psychosis data set. Multiset canonical correlation analysis with joint independent component analysis (mCCA+jICA) is an unsupervised analysis framework that identifies relationships across modalities and decomposes data to reveal maximally independent latent sources of variance (27, 31) (see *supplement*). Briefly, mCCA (39, 40) first aligned the three imaging modalities in order to simplify the correlational structure and maximize inter-subject covariation (28, 30). Next, jICA maximized spatial independence (41) yielding a set of 9 independent components (ICs). Each IC contained a set of linked modalities including maps of cortical thickness (sMRI) and working memory task activation (tfMRI), and a parcel-wise correlation matrix (rsfcMRI). Each IC had a corresponding set of subject-specific weights that reflected the extent to which a given IC comprised the participant’s original data. These weights were then used for statistical analyses to identify brain-behavior relationships and assess group discriminability (see *supplement*).

### Statistical analyses

Statistical analyses were performed in SPSS and MATLAB with multiple comparison correction using FDR (42). For correlations between subject-specific weights and cognitive control performance, partial correlation was used to correct for differences in group means. We assessed group discrimination performance of each IC using multivariate analysis of variance (MANOVA) in which group was used to predict all three imaging weights (one per modality) related to each IC. Significant MANOVA omnibus tests were followed up with planned contrasts.

## Results

### Behavioral and demographics

There were no significant group differences in gender or parental education\SES. Groups differed significantly on age, ethnicity, education, symptoms, and cognitive control performance. SZ had significantly impaired cognitive control performance as compared to HC and BP, with no difference between HC and BP (table 1).

**Table 1:** Demographics, Clinical symptoms, and Behavioral Performance

|  | **Healthy Controls (HC)** | **Bipolar (BP)** | **Schizophrenia (SZ)** | **Omnibus test statistic** | **p-values** |
| --- | --- | --- | --- | --- | --- |
| **Age** | 24.6±3.1 (30) | 26.7±3.0 (27) | 24.9±3.7 (23) | F(2,77)=3.21 | Omnibus p=0.046<br>HC vs SZ p=0.943<br>HC vs BP p=0.049<br>BP vs SZ p=0.145 |
| **Gender** | 14F,16M | 13F,13M | 4F,18M | $\chi^2=6.034$ | Omnibus p=0.197 |
| **Ethnicity (%Caucasian)** | 57% | 80% | 29% | $\chi^2=0.006$ | Omnibus p=0.002<br>HC vs SZ p=0.047<br>HC vs BP p=0.066<br>BP vs SZ p<0.000 |
| **Years of education** | 16.2±2.4 (30) | 15.3±2.3 (26) | 13.0±1.4 (22) | F(2,75)=14.9 | Omnibus p<0.000<br>HC vs SZ p<0.000<br>HC vs BP p=0.264<br>BP vs SZ p=0.001 |
| **Parental level of education / SES proxy** | 2.04±1.3 (27) | 2.28±1.3 (25) | 2.76±1.1 (17) | F(2,66)=1.71 | Omnibus p=0.189 |
| **SAPS** | 0.1±0.7 (30) | 2.2±2.9 (26) | 4.3±3.1 (22) | F(2,75)=19.42 | Omnibus p<0.000<br>HC vs SZ p<0.000<br>HC vs BP p=0.005<br>BP vs SZ p=0.010 |
| **SANS** | 0.8±1.7 (29) | 3.4±3.5 (26) | 6.3±3.0 (22) | F(2,74)=23.69 | Omnibus p<0.000<br>HC vs SZ p<0.000<br>HC vs BP p=0.003<br>BP vs SZ p=0.002 |
| **WERCAP Psychosis history** | 0.90±3.2 (30) | 8.78±7.4 (27) | 31.83±11.59 (23) | F(2,77)=105.76 | Omnibus p<0.000<br>HC vs SZ p<0.000<br>HC vs BP p=0.001<br>BP vs SZ p<0.000 |
| **Cognitive control composite** | 0.44±0.51 (30) | 0.15±0.62 (27) | -0.44±0.97 (23) | F(2,77)=10.31 | Omnibus p<0.000<br>HC vs SZ p<0.000<br>HC vs BP p=0.264<br>BP vs SZ p=0.012 |

Text: mean +/- std (n). N-values vary slightly due to missing data. F=Female; M=Male.
Level of education scale: 1=Graduate professional training; 2=Completed undergraduate; 3=Partial college; 4=High school graduate; 5=Partial high school; 6=Junior high school; 7=less than seven years of education.

### Prediction using a priori ICs

The application of two ICs from (30) (figs. S1-2) generated a set of participant-specific weights corresponding to the extent that these ICs reflected the present dataset. These weights were correlated with the cognitive control composite using partial correlations correcting for group mean differences (table 2, fig. 1 and fig. S3). This demonstrated that patterns contained within tfMRI maps from both *a priori* ICs and the sMRI map from HCP_C1_IC7 were similarly predictive of cognitive control in the psychosis dataset (3/5 modalities significant).

**Table 2:**
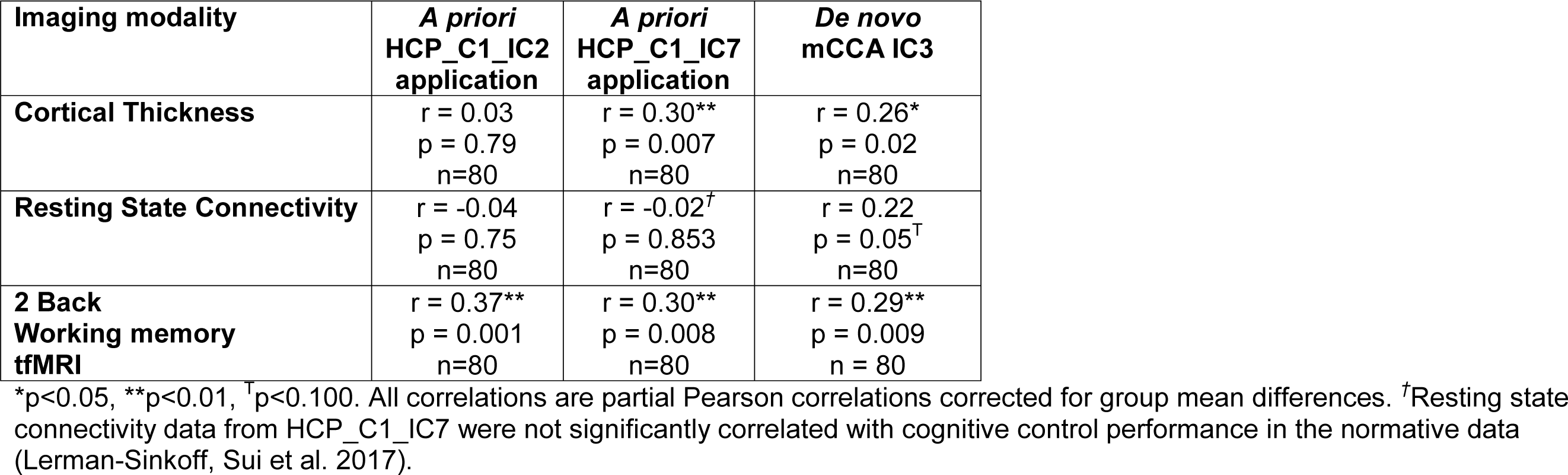
Partial correlation results for application of a priori components and de novo component

**Figure 1:**
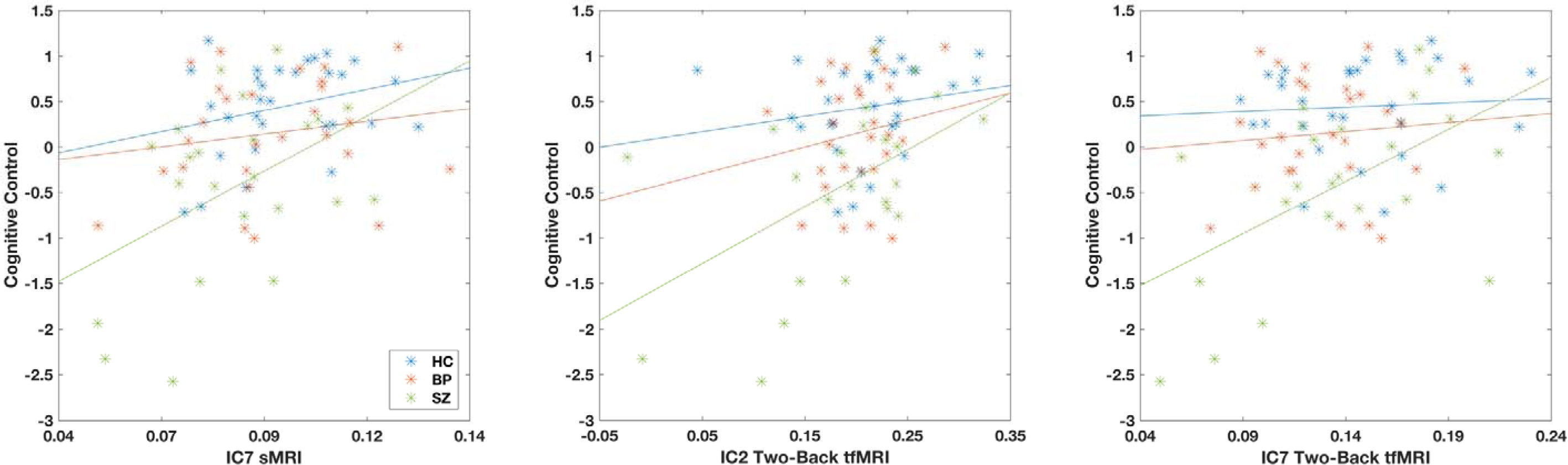
Scatter plots of cognitive control and *a priori* ICs applied to psychosis cohort. Three out of five modalities in HCP_C1_IC2 and HCP_C1_IC7 significantly predicted cognitive control performance in the psychosis cohort. Scatter plots of the other two modalities are available in fig. S3.

**Figure 2:**
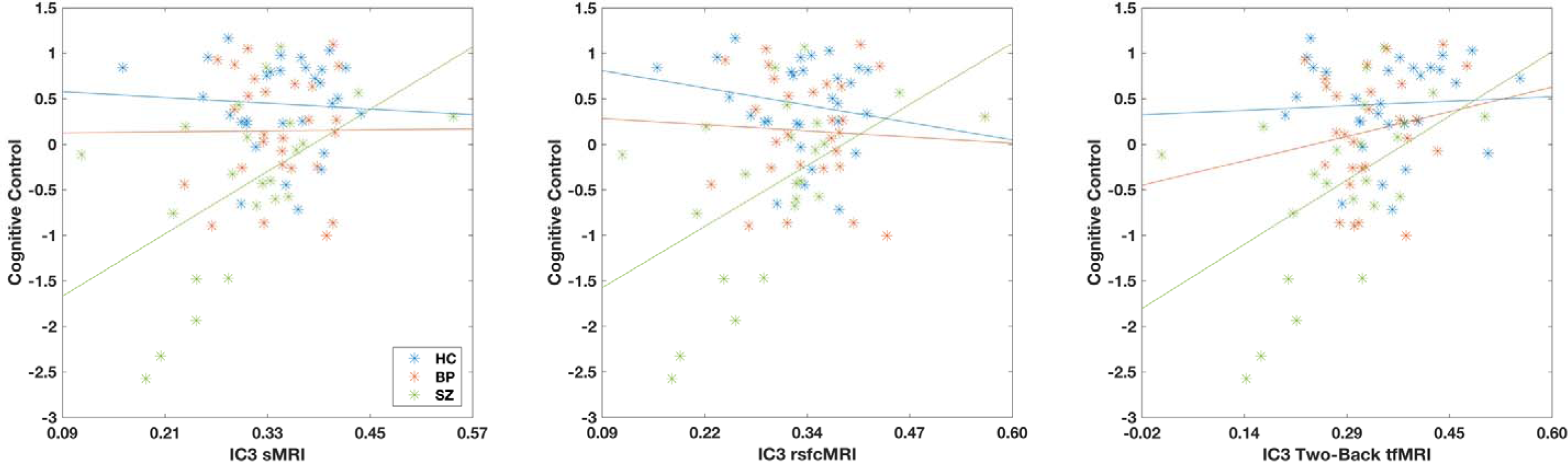
Scatter plots of cognitive control and *de novo* IC3 imaging weights. Partial correlations between *de novo* IC3 imaging weights and cognitive control for all three groups pooled were significant for sMRI and two-back tfMRI modalities (rsfcMRI trended significant). Scatter-plots by group suggested that this may be driven by SZ participants, which was formally tested using regression analyses (tables S6-S8).

### Independent application of mCCA+jICA

mCCA+jICA was also performed *de novo* using the present transdiagnostic cohort. This generated nine ICs and corresponding participant-specific weights. All 27 weights (9 ICs × 3 modalities / IC) were correlated with the cognitive control composite using full Pearson correlation and corrected using FDR. This identified a single component, IC3, that was significantly correlated with cognitive control performance for all three modalities. Follow-up partial correlations correcting for group were performed with FDR correction, indicating that both sMRI and tfMRI still significantly correlated and rsfcMRI trend-level correlated with cognitive control (table 2).

### Spatial distributions of de novo IC3

The modalities in *de novo* IC3 (fig. 3) were visually inspected and bore strong visual resemblances to *a priori* HCP_C1_IC2 (fig. S1). Algorithmic matching using eta^2^ across the full set of *de novo* and *a priori* ICs confirmed visual similarity by pairing IC3 with HCP_C1_IC2 with 61% shared variance across the two ICs (table S1). For sMRI data in IC3 (Fig. 3a), the strongest positive contributing areas were in bilateral anterior and posterior insula, temporal pole, cingulate, frontal superior cortex, and temporal gyrus. Thus, improved cognitive control performance was related to greater thickness in these areas. The strongest negative contributions were in bilateral somatomotor and medial visual cortex, with improved cognitive control performance related to lower thickness in these areas.

For rsfcMRI data in IC3 (Fig. 3b), the strongest positive contributions were concentrated in within-network connectivity, the diagonal of the correlation matrix, with improved cognitive control performance related to stronger within network connectivity. Negative contributions were predominantly concentrated off-diagonal and between the default mode network (DMN) and the task positive networks, including the fronto-parietal, cingulo-opercular, parietal encoding and retrieval, dorsal attention, and ventral attention networks. Thus, improved cognitive control performance was related to stronger negative connections between networks.

**Figure 3a:**
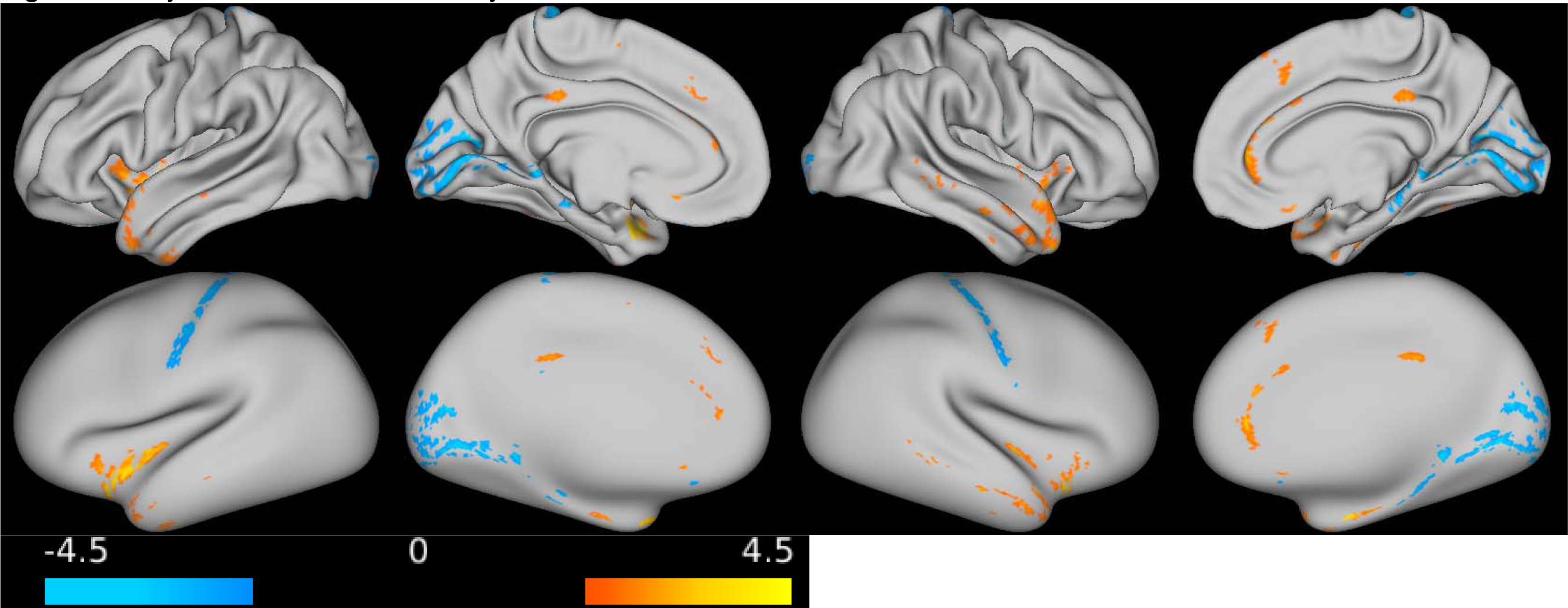
Psychosis IC3 – sMRI modality. Given that mCCA+jICA generates component maps with a value at every vertex/voxel/pairwise-correlation, maps were thresholded at |Z|>2 to simplify interpretation of the spatial pattern of results in de novo IC3 (Sui, Pearlson et al. 2015, Lerman-Sinkoff, Sui et al. 2017). Thus, the maps presented in (Fig, 3a-c, S1, S2) highlight those elements in the map that were strongest relative to all other elements in the given map (unthresholded maps are available in the supplement, figs. S4-S6).

**Figure 3b:**
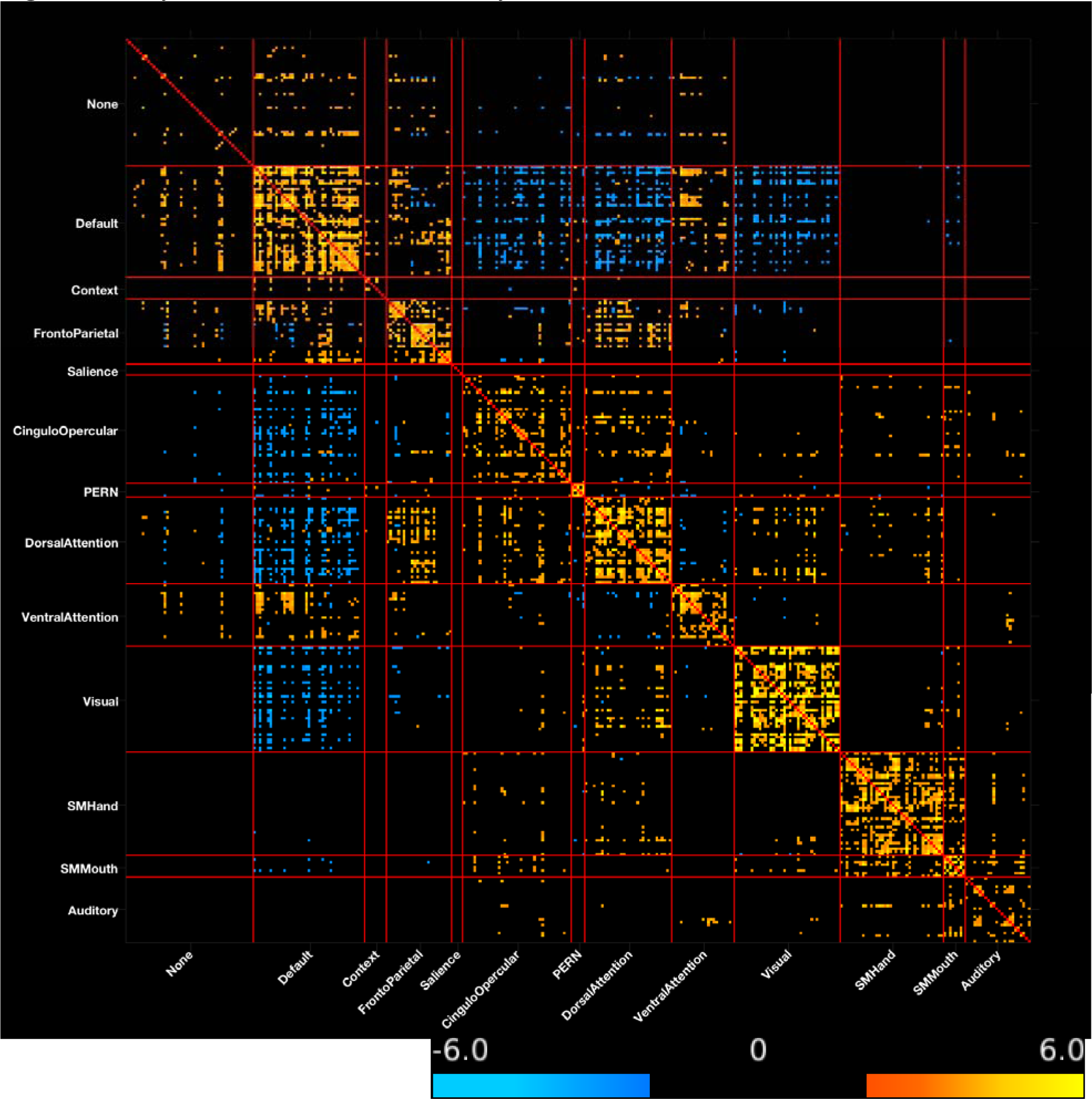
Psychosis IC3 – rsfcMRI modality.

**Figure 3c:**
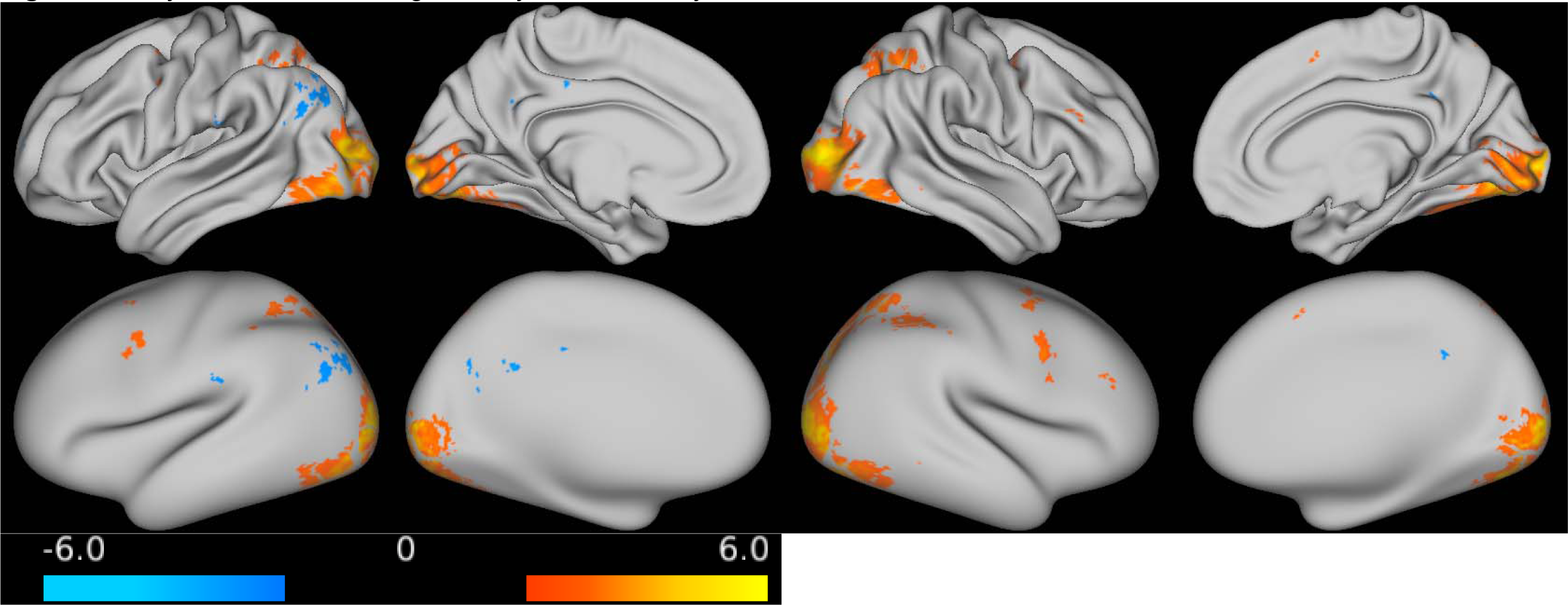
Psychosis IC3 – working memory tfMRI modality.

For tfMRI data in IC3 (Fig. 3c), the strongest positive contributing areas were in bilateral visual cortex, intraparietal sulcus, superior parietal gyrus, precentral sulcus and right middle cingulate, and inferior frontal sulcus, with improved cognitive control performance related to greater activation in these regions. The strongest negative contributions were seen in bilateral isthmus of the cingulate, left supramarginal gyrus, precuneus, and inferior parietal gyrus. Thus, improved cognitive control performance was related to lower activation in these regions.

### Group differences in correlations

Visual examination of scatter plots of participant-specific weights and cognitive control performance suggested that correlations may be driven by SZ (figs. 1 and 2, and fig. S3). Regression analyses were performed for each modality to formally test this observation (tables S3-S8). For the *a priori* data, we tested the three modalities that significantly predicted cognitive control. The only significant interaction effect in *a priori* data was observed in HCP_C1_IC7 tfMRI (table S3), driven by a significant correlation between participant-specific weights and cognitive control for SZ, but not for HC, BP, or HC and BP pooled. Thus, SZ were responsible for the significant correlation in HCP_C1_IC7 tfMRI, but not the other two *a priori* modalities. For *de novo* data, there were significant interaction effects in all three modalities in IC3 (tables S6-S8). Similar to *a priori* HCP_C1_IC7 tfMRI, effects were driven by significant correlations between participant-specific weights and cognitive control for SZ, but not for HC or BP.

**Table 3:** IC3 group-discrimination MANOVA results

|  | Item | Value | Test Statistic | p-values |
| --- | --- | --- | --- | --- |
| **Omnibus tests** | Pillai's Trace | 0.219 | F(6,154)=3.15 | p=0.006 |
|  | Wilk's Lambda | 0.78 | F(6,152)=3.21 | p=0.005 |
| **Between Subjects Effects** | sMRI | n/a | F(2,78)=2.52 | p=0.087 |
|  | rsfcMRI | n/a | F(2,78)=1.91 | p=0.156 |
|  | tfMRI | n/a | F(2,78)=4.59 | p=0.013 |
| **Post-hoc tests** | **Item** | **Mean Difference** | **Standard Error** | **p-values** |
|  | sMRI: HC - BP | 0.001 | 0.018 | p=0.936 |
|  | sMRI: HC - SCZ | 0.038 | 0.018 | p=0.044 |
|  | sMRI: BP - SCZ | 0.036 | 0.019 | p=0.060 |
|  | rsfcMRI: HC - BP | -0.008 | 0.019 | p=0.679 |
|  | rsfcMRI: HC - SCZ | 0.030 | 0.020 | p=0.132 |
|  | rsfcMRI: BP - SCZ | 0.038 | 0.020 | p=0.066 |
|  | tfMRI: HC - BP | 0.025 | 0.022 | p=0.255 |
|  | tfMRI: HC - SCZ | 0.070 | 0.023 | p=0.003 |
|  | tfMRI: BP - SCZ | 0.045 | 0.024 | p=0.066 |

### Group-discrimination

We also performed MANOVAs to assess group-discriminative performance of each *de novo* IC wherein group membership was used to simultaneously predict participant-specific weights for all three modalities in the given IC. Omnibus MANOVA results were significant with FDR correction only for IC3, the only component significantly related to cognitive control (table 3); all other ICs were not group-discriminative (table S2). Post-hoc tests between groups were significant for group differences between HC and SZ for both sMRI and tfMRI. There were trend-level significant differences between BP and SZ for all three modalities, and no differences between HC and BP. Although groups did not significantly differ on parental SES, inclusion of parental SES as a covariate improved the model such that there were trend or significant effects for all modalities (table S9).

## Discussion

The goal of the present study was to examine transdiagnostic multimodal neural alterations in psychosis related to cognitive control. We first used cognitive control related ICs that were identified in the HCP normative population to predict results in our transdiagnostic psychosis cohort. *A priori* working memory tfMRI maps and one sMRI map significantly predicted cognitive control in the psychosis cohort. We then performed mCCA+jICA *de novo* with psychosis data and identified a single IC that significantly correlated with cognitive control performance. This IC exhibited similar spatial distribution to an *a priori* IC and was the sole group-discriminative IC from the *de novo* mCCA+jICA results. Importantly, as described below, our multi-modal analyses illustrate the ways in which there are both deficits in the same regions/networks that cut across modalities, as well as deficits that are modality specific, though contributing to joint prediction of cognitive control. Such results help to link findings in individual modalities of neural structure/function into an associated pattern that is correlated with a central domain of cognitive impairment in psychosis.

Our previous multimodal work studying cognitive control in healthy individuals identified the strongest neural contributions in two distinct patterns: HCP_C1_IC2 contained predominantly posterior cortical contributions in the tfMRI data from the fronto-parietal (FP), dorsal attention (DA), and visual networks. HCP_C1_IC7 contained predominantly dorsolateral and medial PFC, IPL, and IPS contributions in tfMRI data from the FP and DA networks. Importantly, these normative tfMRI patterns were also predictive of cognitive control in our psychosis participants, as well as the sMRI component from HCP_C1_IC7. In this sMRI component, the strongest positively contributing clusters were located in the CO, salience, and ventral attention networks and strongest negative contributing clusters were located in the default mode and visual networks. While these results predominantly include regions in tfMRI data hypothesized to support rapid-timescale cognitive control functionality (43), the sMRI data includes some regions hypothesized to support stable-timescale control (43). Together, these *a priori* patterns were predictive of cognitive control in the general population as well as individuals with varying levels of psychosis and may provide clues towards localization of deficits in psychosis.

For tfMRI data, *a priori* HCP_C1_IC2 and *de novo* IC3 exhibited very similar patterns. We previously postulated that strong visual contributions may be due to top-down modulation of visual regions, especially given that the working memory tfMRI task was highly visually demanding (44). The independent identification of this pattern *de novo* may provide intriguing clues towards the source of cognitive control dysfunction in SZ. A number of reports have identified visual system dysfunction in SZ (45–48), though the literature is more mixed for BP (49–54). Theories generated from these lines of research postulate that alterations in visual system functionality leads to aberrant integration of information in higher order cortical areas and lead to dysfunction in cognitive control (45, 48, 55). Thus, the present work could be seen as consistent with the hypothesis that impairments in the function and/or structure of visual cortex may disrupt higher order processing leading to deficits in cognitive control. In the present data, SZ participants who exhibited greater visual region contributions in IC3 tfMRI data had significantly better cognitive control performance. Importantly, the data for HC and BP trended in the same direction as SZ, though this trend was not significant (fig. 2). While the present work is not a direct assessment of the relationship between visual system integrity and cognition, the findings are consistent with these models of psychosis in SZ and thus warrant further study.

For sMRI data, both *a priori* HCP_C1_IC2 and *de novo* IC3 exhibited relationships between greater cortical thickness and cognitive control performance in the insula, medial PFC, and cingulate. However, *de novo* IC3 also included negative contributions in somatomotor and medial visual cortex. These findings are consistent with the extant cognitive control related unimodal sMRI literature. For example, dichotomizing BP and SZ into high and low executive function groups identified greater cortical thickness in right pre- and post-central gyri in the high functioning group as compared to the low functioning group (18). While there are few reports of relationships between cognitive control and visual cortex thickness in psychosis, there is evidence for differential cortical thickness alterations between BP and SZ in early retinotopic cortex (56). This region showed a relationship with cognitive control in the present data for SZ participants. More anteriorly, cortical thickness alterations have been identified transdiagnostically in the anterior cingulate and insula but relationships with cognitive control in psychosis were not tested (17). Interestingly, these same anterior regions were present in both our *a priori* and *de novo* ICs and relationships with cognitive control were significant for all three groups in *a priori* data. Given that *de novo* data were only significant for SZ, this suggests the possibility that anterior cortical thickness contributions to cognitive control may be transdiagnostic and somatomotor and visual negative contributions may be unique to SZ.

For rsfcMRI data, *a priori* HCP_C1_IC2 was not predictive of cognitive control in psychosis. Further, *de novo* rsfcMRI data from IC3 only trend-level correlated with cognitive control. Similar to rsfcMRI data from HCP_C1_IC2, *de novo* rsfcMRI data in IC3 exhibited a modular network pattern comprised of high within-network connectivity and anti-correlated activity between the task-positive and task-negative networks and resembled a canonical resting state matrix. Thus, we draw the same general conclusion as in our prior work, namely that there is some evidence that greater presence of canonical resting state networks in *de novo* data may be associated with improved cognitive control. While relationships between ICs and cognitive control in *a priori* analyses were consistent across groups (excluding HCP_C1_IC7 tfMRI), relationships across groups for *de novo* data were less so. Significant correlations between *de novo* IC3 and cognitive control were driven by SZ, which may be due to a variety of factors. First, compared to BP and HC, SZ were significantly worse on all metrics including IC weights, cognitive control performance, and clinical variables. Importantly, there was significant inhomogeneity of variance such that SZ had significantly more variance in cognitive control than HC and numerically greater than BP, which may have hampered our ability to identify relationships with cognitive control in the other groups. Further, BP inclusion was not limited to individuals with psychosis and BP endorsed significantly less history of psychotic symptoms (table 1). Importantly, the literature suggests variability in cognitive performance in BP (57) with poorer working memory and executive function performance in psychotic BP. Indeed, dichotomizing our BP participants by psychosis severity revealed that low-psychosis BP performed significantly better than SZ and high-psychosis BP exhibited performance indistinguishable from SZ (table S10). Finally, modalities that showed significant between-subjects effects exhibited a consistent pattern of mean imaging weights (HC > BP > SZ). Thus, future work with larger BP samples with psychosis are needed to better determine similarity to schizophrenia in multimodal correlates of cognitive control.

There are several limitations to the present study. First, *de novo* mCCA+jICA was a data-driven explorative analysis and must be interpreted with caution. However, given similarity between *a priori* ICs and the present results, the differences across datasets provide intriguing possibilities for further hypothesis-driven study. Second, the present cognitive control metric was not identical to the HCP metric due to study design differences, potentially limiting our ability to detect all of the same relationships identified in our prior work. Third, while IC3 was group discriminative overall, we were unable to detect significant post-hoc differences between HC and BP, though differences between HC and SZ were significant and differences between SZ and BP were trend-level. This may be due to limited sample size, medications, or a relatively unimpaired BP group.

In conclusion, the present study employed multimodal methods to examine transdiagnostic alterations in cognitive control in psychosis. Two *a priori* ICs from the normative population significantly predicted cognitive control in psychosis for three of five modalities tested. *De novo* mCCA+jICA identified a group-discriminative IC that significantly correlated with cognitive control for sMRI and tfMRI and trend-level with rsfcMRI. *De novo* analyses suggest joint associations between cognitive control and tfMRI contributions from the posterior frontoparietal, dorsal attention, and visual networks, sMRI contributions from the insula, medial PFC, cingulate, somatomotor and visual regions, and possibly canonical resting-state network organization and point to unique contributions in SZ from somatomotor and visual cortices. However, significant *de novo* effects were driven by SZ with little evidence for effects in BP. Given psychotic symptom heterogeneity in BP, results suggest that shared symptomatology, e.g. psychosis, may be key to identification of transdiagnostic relationships with cognitive control. Together, these results identified significant and replicable relationships across modalities and the psychosis spectrum, providing novel targets across modalities of neural structure and function for future research.

## Acknowledgements

DBLS was supported by NIH MSTP training grants 5T32GM007200-38, 5T32GM007200-39; Interdisciplinary Training in Cognitive, Computational and Systems Neuroscience 5 T32 NS073547-05; the McDonnell Center for Systems Neuroscience; and NIH fellowship F30MH109294. DTM was supported by NIMH grant R01 MH104414; Taylor Family Institute, Dept. of Psychiatry; and the Center for Brain Research on Mood Disorders, Dept. of Psychiatry, Washington University Medical School. VDC was supported by NIH grants 2R01EB006841, 2R01EB005846, P20GM103472, and NSF grant 1539067. Computations were performed using the facilities of the Washington University Center for High Performance Computing, which were partially funded by NIH grants 1S10RR022984-01A1 and 1S10OD018091-01. Figures S1 and S2 are reprinted from Neuroimage, Volume 163, Authors Dov Lerman-Sinkoff, Jing Sui, Srinivas Rachakonda, Sridhar Kandala, Vince Calhoun, Deanna Barch, Multimodal neural correlates of cognitive control in the Human Connectome Project, 41-54, Copyright 2017, with permission from Elsevier. The content of this report is solely the responsibility of the authors and does not necessarily represent the views of the funding agencies. This manuscript is intended for submission to bioRxiv and has not been peer reviewed.

