## Supplementary Materials for "Transdiagnostic multimodal neuroimaging in psychosis: structural, resting-state, and task MRI correlates of cognitive control"

**Supplementary Methods**:

*Participants*

N=31 healthy controls (HC), n=27 persons with bipolar disorder (BP), and n=23 persons with schizophrenia spectrum disorders (SZ) were included in the study. One HC participant contributed imaging data but lacked behavioral measures, leaving n=30 HC and a total of N=80 participants used for correlations between imaging and cognitive control performance.

*Behavioral assessment*

Three individual measures were used to generate the composite metric of cognitive control: (1) In-scanner working memory task – accuracy in the 2-back condition: participants performed the in-scanner working memory task used in the HCP (1). This task consisted of a block design in which participants performed both 0-back and 2-back blocks with one of four possible stimuli: faces, places, body parts, and tools. Participants were asked to respond when the presented stimulus was the same as the stimulus displayed two stimuli prior, and 20-30% of stimuli were lures (1-back and 3-back) to help ensure the use of active memory techniques rather than stimulus familiarity. (2) Out of scanner working memory task – accuracy in the 2-back condition: participants performed the Penn Computerized Neuropsychological Testing battery (2), which included a N-back task with letters as stimuli and three blocks of 0-back, 1-back, and 2-back conditions. 2-back accuracy scores were used and computed as the number of true positive responses minus the number of false positive responses in the 2-back block. (3) Penn Progressive matrices – total number correct. This was a shortened version of the classic Raven's Progressive Matrices task wherein participants were presented with a texture in which a section was removed. Participants were asked to choose the pattern that best completed the texture from a set of possible options. Given that not all participants had full data (i.e.: some participants lacked one or two of the aforementioned measures), the composite measure was generated by first Z-scoring the individual metrics and then taking the mean of the available metrics for any given participant.

*Neuroimaging collection and pre-processing*

Imaging data were collected at Washington University in St. Louis using the customized Siemens “Connectome” Skyra scanner equipped with 100 mT/m gradient coils and a 32-channel head coil. BOLD contrast images were collected using gradient-echo echo-planar imaging accelerated with an 8X multiband sequence. Resting state data were collected with eyes open and crosshair fixation (3) over two days with two 864 second scans per day (in both L->R and R->L phase encoding directions). The working memory task data were collected in two scans (both phase encoding directions, as above) with 301 seconds per scan (1).

Structural scans were collected and processed with the HCP minimal preprocessing pipelines (4). Briefly, T1 and T2 images were processed in three sequential stages. The first pipeline performed correction of gradient nonlinearity-induced distortions, alignment of subject scans to MNI coordinate space, removal of readout-distortions, correction of intensity inhomogeneities, and finally alignment to MNI atlas space. The second pipeline processed data using an HCP-customized version of Freesurfer and generated participant-specific segmentations and parcellations. Finally, the third pipeline converted Freesurfer output into NIFTI, CIFTI, and GIFTI formats and registered data to several surface meshes including the “32k_fs_LR” mesh used in the present study.

Working memory task and resting-state data were also processed with the HCP pipelines (4). Briefly, collected data in both phase encoding directions were processed by performing gradient unwarping, motion correction, EPI field distortion correction, registration to T1w data, registration into MNI space, and intensity normalization. Next, the cortical ribbon was projected to the surface and registered to the meshes generated in the structural pipeline in order to align all subjects' data into a standard "grayordinates" space which included cortical data projected to the surface as well as volumetric data for the subcortex and cerebellum. Surface and volume data were smoothed separately in order to achieve a final smoothing of 4mm FWHM. The HCP minimal preprocessing pipeline for resting state data ended here (see below for further post-processing). Working memory task data were then further processed with the HCP FSL pipeline in order to generate subject-specific contrast maps corresponding to activation in the 2-back condition.

Resting state data were further processed to transform vertex\voxel timecourses into parcel-wise correlation matrices as was done in (5). Resting state data were demeaned and detrended within each run and a single regression was used to remove twenty-four motion related parameters (six rotational and translational parameters, their derivatives, and squares), the mean grayordinates timeseries (akin to global signal) (6), and noise components from FSL MELODIC identified by FIX (7, 8). Data were then highpass filtered (cutoff frequency 0.009 Hz), demeaned and detrended, individual runs concatenated, and mean parcel timeseries were extracted using a cortical parcellation (9), cerebellar parcellation (10), and subcortical regions identified by Freesurfer. Functional connectivity matrices were then generated for each participant as the pairwise Pearson correlation between all parcels and the lower-triangle of the symmetric correlation matrix was used for mCCA+jICA analysis to avoid incorporation of redundant data.

*mCCA+jICA parameter selection*

A limitation of the mCCA+jICA model is that there is relatively little guidance as to the optimal number of ICs to select for decomposition of the source data or the number of singular values (SVs) to use during dimensionality reduction (5). To address this, we used the results and methodology described in (5) to guide parameter selection in the present dataset. Given that the overarching goal was to identify domains of variation related to cognitive control that span the spectrum from normative to psychotic function, we chose an identical number of ICs (9 independent components) as was done in the previous study. To determine the optimal number of SVs, we performed an analysis using the Washington University Center for High Performance Computing cluster to sweep through all possible numbers of SVs (one through eighty-one) and used ICASSO (described below) to generate a metric of analysis stability at a given SV. Based on that analysis, we chose a value for SV (76) that maintained a large amount of variance (greater than 98% in all modalities) in the data while still being in the neighborhood of other SVs that generated stable results (high values of analysis stability, termed Iq, described below).

*mCCA+jICA methodology*

The full theory and methodology of mCCA+jICA have been previously published (11-16) and an overview of the methodology is presented here. Similar to (5), an in-house modified version of the FIT toolbox (http://mialab.mrn.org/software/fit) was used to perform mCCA+jICA. The analysis was performed in three stages: dimensionality reduction, multiset canonical correlation analysis (mCCA), and joint independent component analysis (jICA). Of note, group labels (HC, BP, or SZ) were not used in the mCCA+jICA decomposition in order to prevent biasing of results. Data files for each participant corresponding to the three modalities of data (sMRI, rsfcMRI, and tfMRI) were loaded into MATLAB, linearized into vectors, and then concatenated to generate three matrices corresponding to the three modalities of data. Dimensionality reduction was performed using a singular value decomposition of the data to enable computational tractability of downstream analyses. The reduced dimensionality dataset was then used for mCCA analysis.

The goal of the mCCA step is to align the three data modalities in a manner that identifies patterns across the modalities while simplifying the correlational structure and maximizing intersubject covariation (17). This process decomposed modalities into a set of group-level components (modality-specific maps) and a set of corresponding mixing profiles (participant loading parameters upon the components). The mixing profiles described the extent to which a given component was reflective of a given participant's data. Each component contained a set of three maps, one per modality, that were linked and represented the extent to which a given vertex/voxel/parcel-wise correlation contributed to the map relative to all other vertices/voxels/parcel-wise correlations within the map. While the analysis could theoretically stop at this point, components generated by mCCA may not be sufficiently separated due to noise and dependencies inherent to neuroimaging data(18-20).

jICA (14) was then performed to address this incomplete separation by further decomposing the components from mCCA. jICA extends traditional unimodal ICA analysis to multimodal data and generates maximally spatially independent latent sources of variance. The component matrices from mCCA were concatenated along the feature dimension and then analyzed 100 times using the infomax ICA algorithm (21) under the ICASSO framework (described below) (22, 23) in order to ensure analysis stability and reproducibility. Similar to mCCA, this process generated a set of group-level independent component (IC) matrices wherein each IC contained linked maps of the three individual modalities and a corresponding set of participant-specific weights that corresponded to the extent to which that modality and component were reflective of a given participant's data. These weights were then used in statistical analyses to assess the relationship between the neuroimaging findings identified by mCCA+jICA and their relationship to cognitive control performance.

*ICASSO*

ICASSO is an ICA analysis framework that helps ensure the stability and reproducibility of ICA analyses (22, 23). ICASSO runs a given ICA analysis N times (N=100 in the present study) with each analysis started with randomly chosen initial conditions. The resultant ICs from the set of N analyses are then clustered and the single IC within a given cluster that is most similar to all other ICs within its cluster is chosen as the final IC for downstream analyses. The clustering process generates a set of cluster quality metrics, termed Iq, that reflect the compactness of a given cluster and its isolation from all other clusters. The Iq metric varies from 0 (lowest quality) to 1 (highest quality) and is computed as the normalized sum of the magnitude of correlations between ICs within a given cluster minus the normalized sum of the magnitude of correlations of ICs within the cluster to ICs outside of the cluster (23). Thus, for a given ICASSO analysis, 9 Iq values were generated, one for each component. These values were then used as described above to select the optimal number of SVs for mCCA+jICA parameter selection. Further, the nine ICs selected by ICASSO in the SV=76 analysis were used in downstream analyses to examine their relationship to cognitive control.

*Relationship to a-priori multimodal normative correlates of cognitive control:*

We performed two analyses to assess the relationship between ICs related to cognitive control that were generated using data from healthy community participants (5) and the present dataset. We first wished to determine whether the ICs that were replicably identified and significantly related to cognitive control in a healthy community sample were also predictive of cognitive control in the present dataset. To do so, we generated participant-specific weightings by multiplying the source data from the present dataset by the pseudoinverse of the ICs determined in (5). These weights were then correlated with the cognitive control metric using partial correlation controlling for group. Second, we also wished to algorithmically match the ICs in (5) to the ICs identified in the de-novo application of mCCA+jICA to the present dataset. To do so, we implemented an eta$^2$ similarity function (24) to assess the extent to which variance in one IC accounted for variance in another IC. This method was chosen in lieu of other distance metrics, such as Pearson correlation, as it accounts for both magnitude and covariance. As was done in (5), we computed a matrix of eta$^2$ values of all possible pairings of ICs in the two datasets using absolute valued ICs to account for sign flipping which may occur due to sign conventions in the mCCA+jICA model. Pairings were made in a 1:1 fashion iteratively without replacement by selecting the highest available value of eta$^2$ in the matrix. Matched ICs were then visually examined to assess the performance of the matching algorithm.

*Software Versions*

- Statistical Analyses – SPSS V25 (IBM, Armonk, NY) and MATLAB R2017a (The Mathworks, Natick, MA).
- mCCA+jICA Analysis – MATLAB R2015a (The Mathworks, Natick, MA).
- Visualization of results - MATLAB R2017a (The Mathworks, Natick, MA) and Connectome Workbench v1.2.3 (Van Essen Lab, St. Louis, MO) (25).
- FDR – fdr_bh, publically available at (http://www.mathworks.com/matlabcentral/fileexchange/27418).
- HCP Pipelines v3.17.0
- FSL v5.0.9
- FIX v1.065, using the HCP training dataset
