## Supplementary Materials for "Transdiagnostic multimodal neuroimaging in psychosis: structural, resting-state, and task MRI correlates of cognitive control"

Figure S1: Reproduction of HCP_C1_IC2

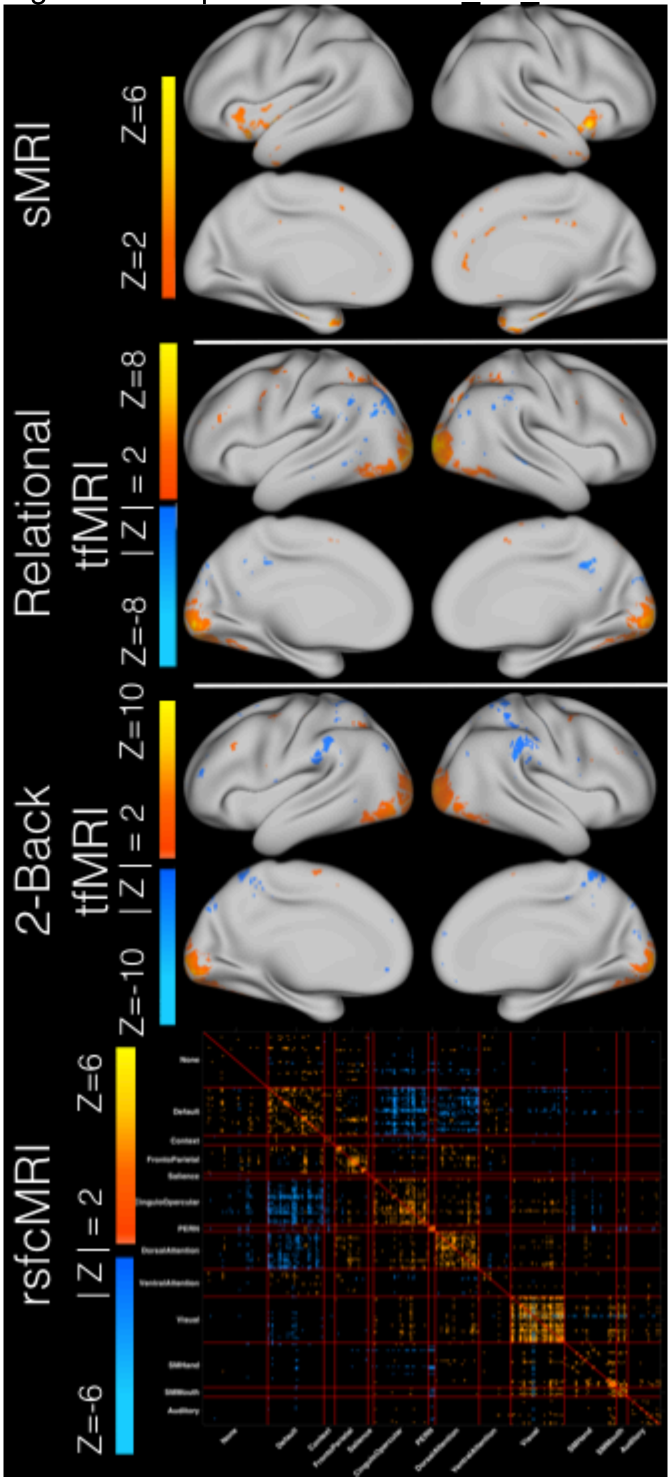

Figure Caption**:** All images show Z-scores of the IC spatial maps for a given modalities' data and are thresholded at |Z| > 2. Note that relational tfMRI was not available for the present study.

Figure S2: Reproduction of HCP_C1_IC7

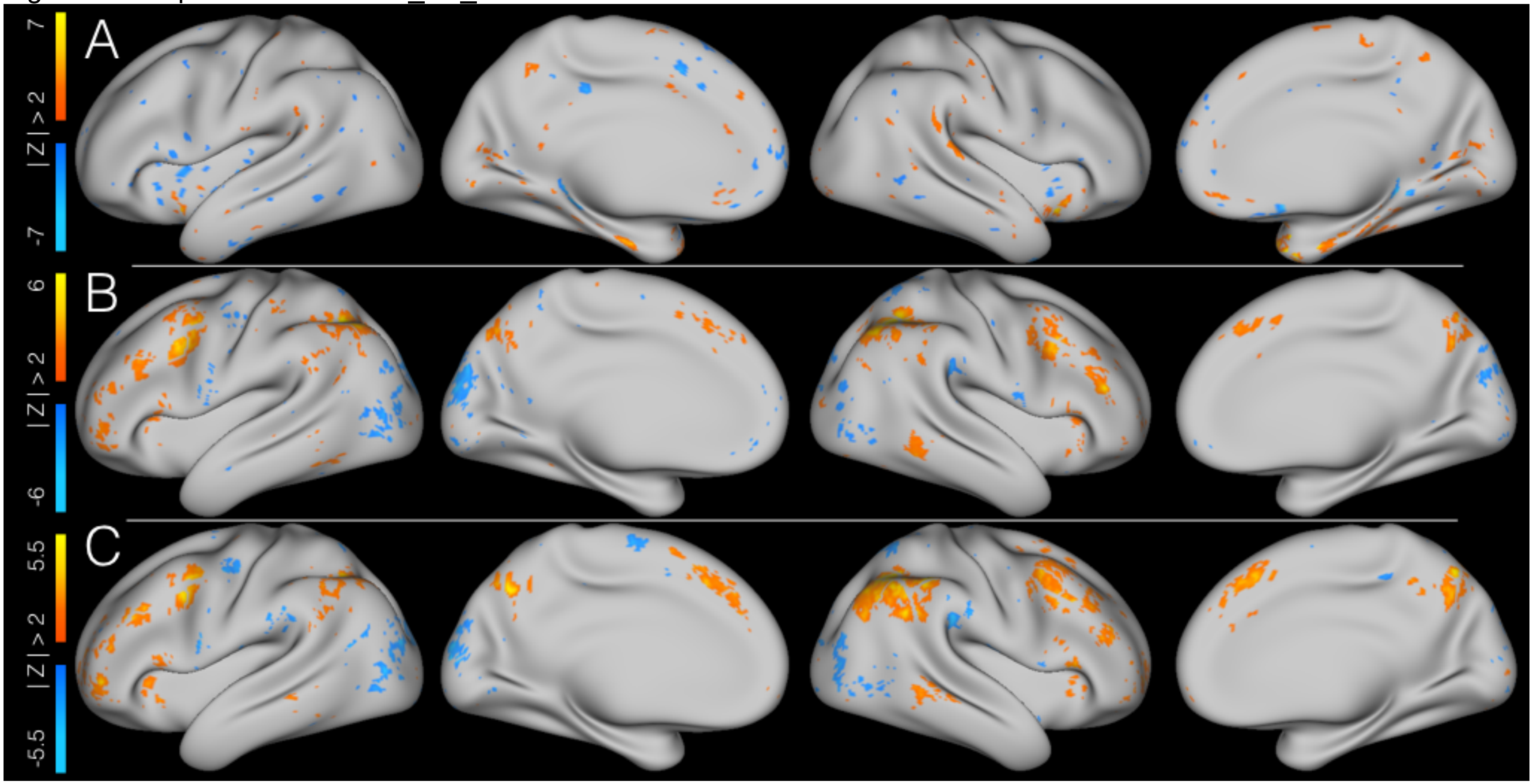

Figure Caption: All images show Z-scores of the IC spatial maps for a given modalities' data and are thresholded at |Z| > 2. A = cortical thickness; B = relational tfMRI (not included in the present study); C = 2-back tfMRI.

Figure S3: Scatter plots of cognitive control and *a priori* ICs applied to psychosis cohort for modalities that did not significantly correlate with cognitive control performance

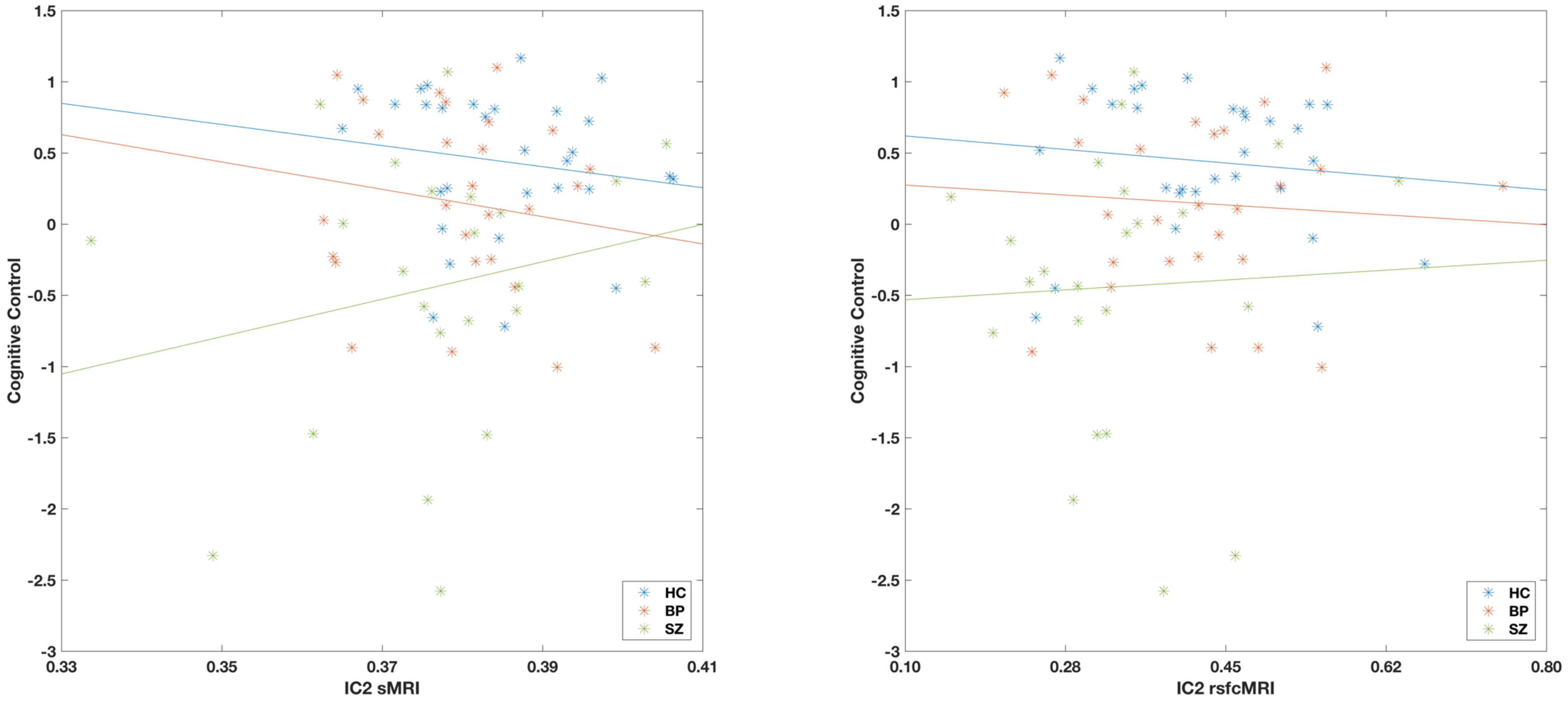

Caption: Two of five *a priori* ICs did not significantly predict cognitive control performance in psychosis.

**Fig S4:** Unthresholded sMRI map from IC3

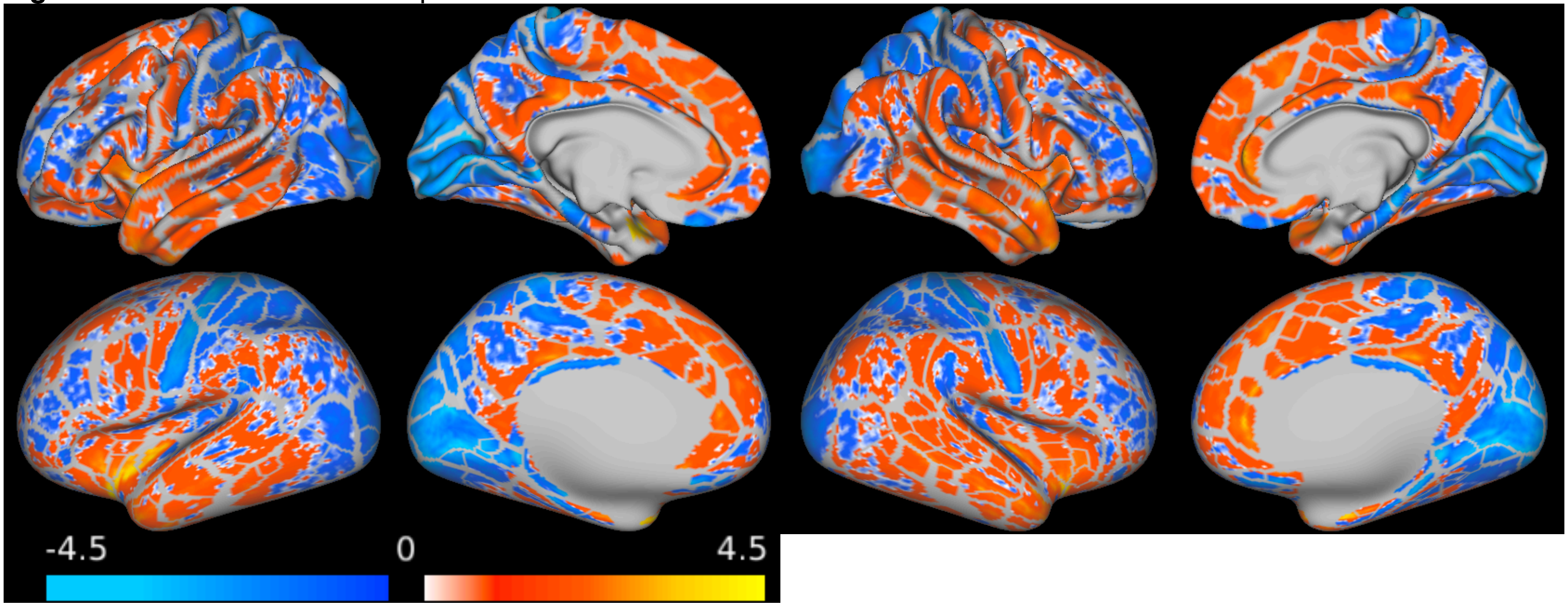

**Fig S5:** Unthresholded rsfcMRI map from IC3

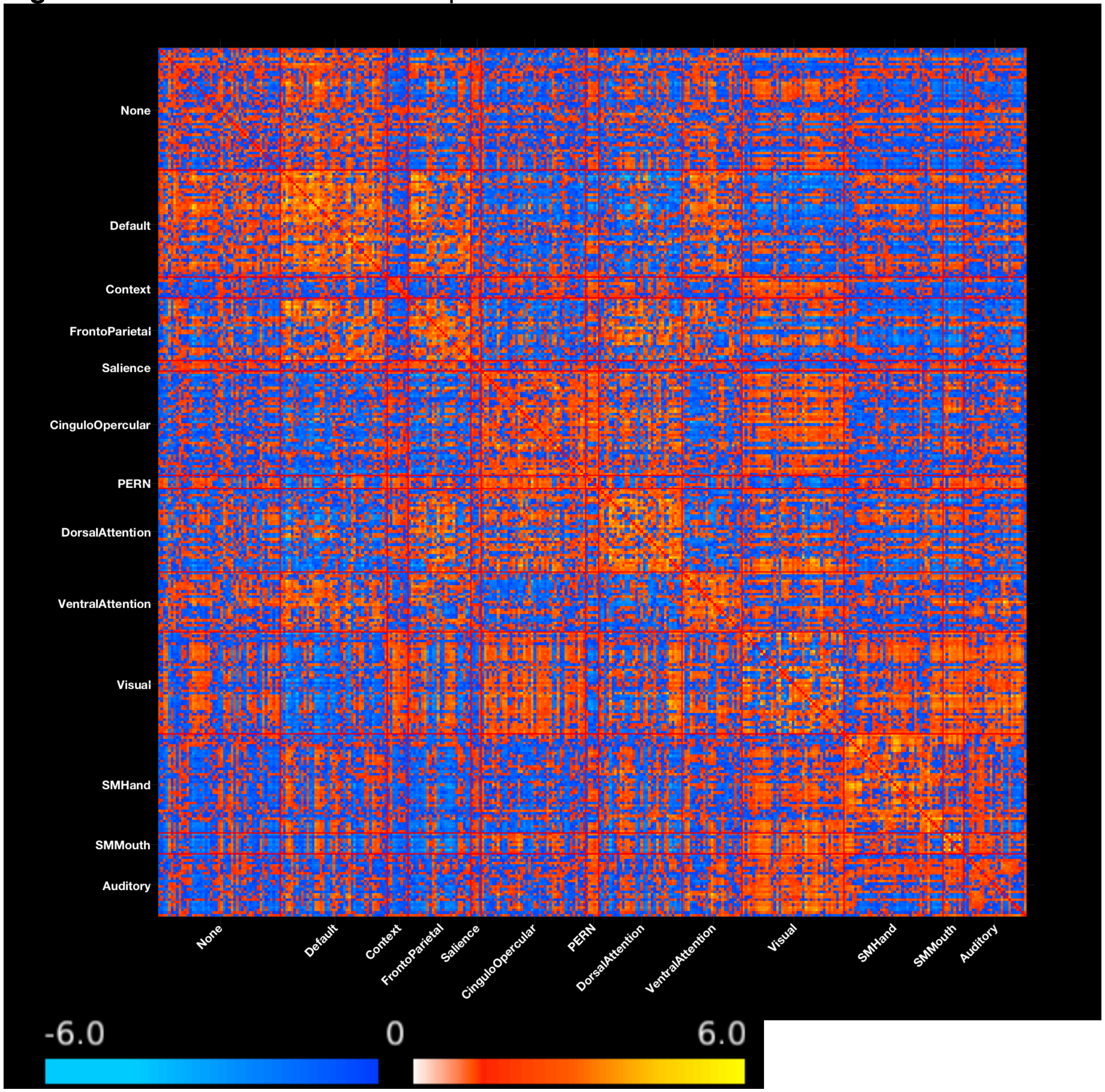

**Fig S6:** Unthresholded working memory tfMRI from IC3

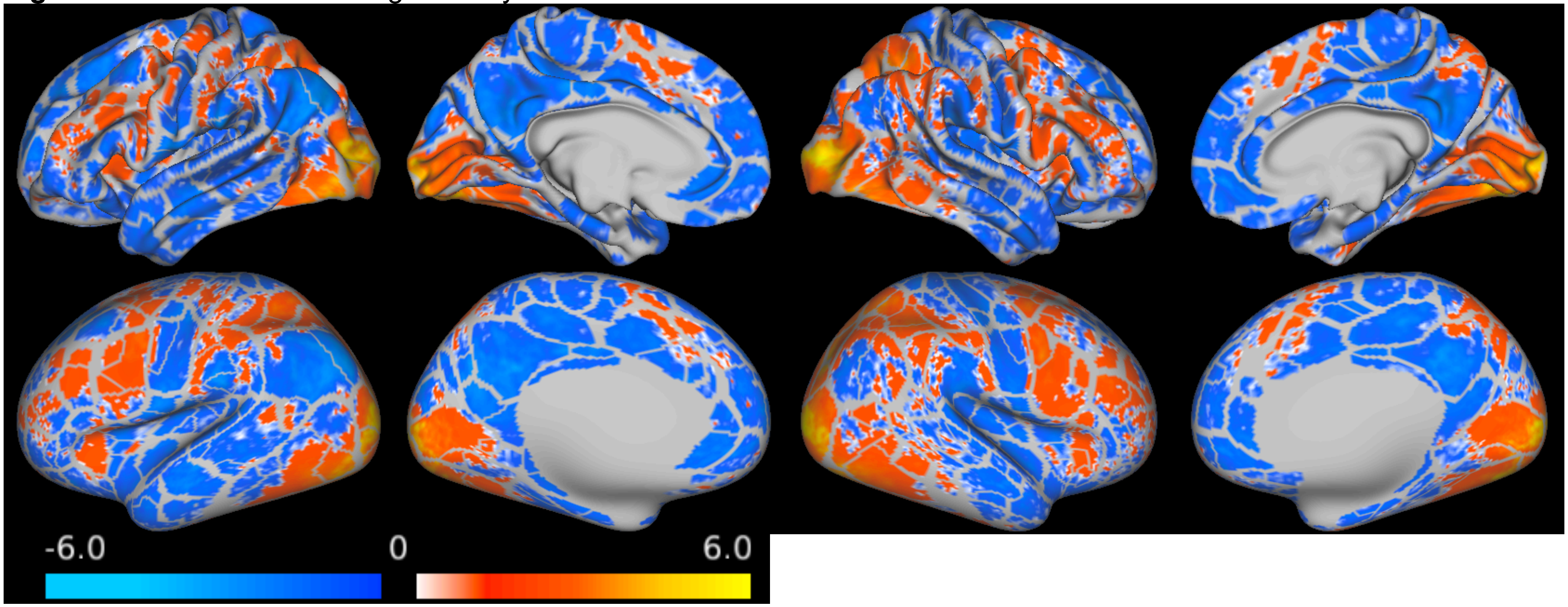

Table S1: Eta$^2$ table

Psychosis cohort (columns); HCP Cohort 1 (rows)

| | IC1 | IC2 | IC3 | IC4 | IC5 | IC6 | IC7 | IC8 | IC9 |
|---|---|---|---|---|---|---|---|---|---|
| IC1 | 0.312 | **0.283** | 0.351 | 0.298 | 0.306 | 0.305 | 0.305 | 0.312 | 0.313 |
| IC2 | 0.538 | 0.528 | **0.609** | 0.515 | 0.557 | 0.533 | 0.529 | 0.542 | 0.568 |
| IC3 | 0.498 | 0.502 | 0.540 | 0.497 | 0.514 | 0.504 | 0.501 | 0.528 | **0.516** |
| IC4 | 0.532 | 0.504 | 0.554 | 0.483 | 0.522 | 0.513 | 0.515 | **0.547** | 0.539 |
| IC5 | 0.318 | 0.295 | 0.361 | **0.309** | 0.316 | 0.312 | 0.315 | 0.324 | 0.323 |
| IC6 | 0.333 | 0.308 | 0.372 | 0.321 | 0.328 | **0.327** | 0.328 | 0.337 | 0.336 |
| IC7 | 0.481 | 0.469 | 0.504 | 0.487 | 0.486 | 0.487 | **0.505** | 0.498 | 0.485 |
| IC8 | **0.350** | 0.325 | 0.391 | 0.344 | 0.348 | 0.347 | 0.349 | 0.36 | 0.354 |
| IC9 | 0.331 | 0.307 | 0.372 | 0.325 | **0.329** | 0.326 | 0.327 | 0.336 | 0.335 |

Table S2: MANOVA tests for Group Discrimination for all *de novo* ICs

| IC | Pillai's Trace Omnibus | Pillai p-values | Wilk's Lambda Omnibus | Wilks Lambda p-values |
|---|---|---|---|---|
| IC1 | F(6,154) = 0.972 | p = 0.446 | F(6,152) = 0.970 | p = 0.448 |
| IC2 | F(6,154) = 1.040 | p = 0.402 | F(6,152) = 1.043 | p = 0.400 |
| IC3 | F(6,154) = 3.153 | p = 0.006 | F(6,152) = 3.214 | p = 0.005 |
| IC4 | F(6,154) = 1.752 | p = 0.113 | F(6,152) = 1.776 | p = 0.107 |
| IC5 | F(6,154) = 1.818 | p = 0.099 | F(6,152) = 1.838 | p = 0.095 |
| IC6 | F(6,154) = 2.323 | p = 0.036 | F(6,152) = 2.373 | p = 0.032 |
| IC7 | F(6,154) = 2.068 | p = 0.060 | F(6,152) = 2.087 | p = 0.058 |
| IC8 | F(6,154) = 1.491 | p = 0.185 | F(6,152) = 1.513 | p = 0.177 |
| IC9 | F(6,154) = 2.272 | p = 0.040 | F(6,152) = 2.280 | p = 0.039 |

Caption: IC3 was the sole group discriminative IC of all nine ICs generated by *de novo* mCCA+jICA

Table S3: *a priori* HCP_C1_IC7 sMRI regression

| Item | Standardized Betas | Test Statistic | p-values |
|---|---|---|---|
| **Omnibus** | n/a | $F(5,74)=6.788$ | $p<0.000$ |
| **sMRI 7 Weights** | $\beta=-0.339$ | $t=-2.45$ | $p=0.006$ |
| **BP Group Membership** | $\beta=0.040$ | $t=0.65$ | $p=0.948$ |
| **SCZ Group Membership** | $\beta=-1.157$ | $t=-1.932$ | $p=0.057$ |
| **sMRI7 * BP interaction** | $\beta=0.203$ | $t=0.341$ | $p=0.734$ |
| **sMRI7 * SCZ interaction** | $\beta=-0.770$ | $t=-1.294$ | $p=0.200$ |

Table S4: *a priori* HCP_C1_IC2 working memory tfMRI regression

| Item | Standardized Betas | Test Statistic | p-values |
|---|---|---|---|
| **Omnibus** | n/a | $F(5,74)=7.700$ | $p<0.000$ |
| **tfMRI 2 Weights** | $\beta=0.350$ | $t=2.278$ | $p=0.026$ |
| **BP Group Membership** | $\beta=-0.322$ | $t=-0.595$ | $p=0.554$ |
| **SCZ Group Membership** | $\beta=-0.977$ | $t=-2.829$ | $p=0.006$ |
| **tfMRI2 * BP interaction** | $\beta=0.163$ | $t=0.298$ | $p=0.766$ |
| **tfMRI2 * SCZ interaction** | $\beta=0.548$ | $t=1.638$ | $p=0.106$ |

Table S5: *a priori* HCP_C1_IC7 working memory tfMRI regression

| Item | Standardized Betas | Test Statistic | p-values |
|---|---|---|---|
| **Omnibus** | n/a | F(5,74)=7.561 | p<0.000 |
| **tfMRI 2 Weights** | β=-0.321 | t=-2.461 | p=0.016 |
| **BP Group Membership** | β=-0.245 | t=-0.527 | p=0.600 |
| **SCZ Group Membership** | β=-1.312 | t=-3.459 | p=0.001 |
| **tfMRI2 * BP interaction** | β=-0.083 | t=-0.177 | p=0.860 |
| **tfMRI2 * SCZ interaction** | β=-0.856 | t=-2.320 | p=0.023 |

Table S6: *de novo* IC3 sMRI regression

| Item | Standardized Betas | Test Statistic | p-values |
|---|---|---|---|
| **Omnibus** | n/a | F(5,74)=7.267 | p<0.000 |
| **sMRI 3 Weights** | β=0.253 | t=2.278 | p=0.060 |
| **BP Group Membership** | β=-0.305 | t=-0.595 | p=0.658 |
| **SCZ Group Membership** | β=-1.616 | t=-2.829 | p=0.002 |
| **sMRI3 * BP interaction** | β=0.126 | t=0.298 | p=0.853 |
| **sMRI3 * SCZ interaction** | β=1.189 | t=1.638 | p=0.0188 |

Table S7: *de novo* IC3 rsfcMRI regression

| Item | Standardized Betas | Test Statistic | p-values |
|---|---|---|---|
| **Omnibus** | n/a | F(5,74)=7.351 | p<0.000 |
| **rsfcMRI 3 Weights** | β=-0.218 | t=-1.737 | p=0.07 |
| **BP Group Membership** | β=-0.370 | t=-0.581 | p=0.563 |
| **SCZ Group Membership** | β=-1.730 | t=-3.56 | p=0.001 |
| **rsfcMRI3 * BP interaction** | β=-0.197 | t=-0.313 | p=0.755 |
| **rsfcMRI3 * SCZ interaction** | β=-1.294 | t=-2.698 | p=0.009 |

Table S8: *de novo* IC3 working memory tfMRI regression

| Item | Standardized Betas | Test Statistic | p-values |
|---|---|---|---|
| **Omnibus** | n/a | F(5,74)=7.052 | p<0.000 |
| **tfMRI 3 Weights** | β=0.359 | t=2.41 | p=0.015 |
| **BP Group Membership** | β=-0.458 | t=-0.848 | p=0.399 |
| **SCZ Group Membership** | β=-1.20 | t=-3.126 | p=0.003 |
| **tfMRI3 * BP interaction** | β=0.294 | t=0.552 | p=0.583 |
| **tfMRI3 * SCZ interaction** | β=0.808 | t=2.167 | p=0.033 |

**Table S9: IC3 group-discrimination MANOVA results including SES covariate**

| | Item | Value | Test Statistic | p-values |
|---|---|---|---|---|
| **Omnibus tests for Group** | Pillai's Trace | 0.219 | F(6,154)=3.153 | p=0.006 |
| | Wilk's Lambda | 0.78 | F(6,152)=3.214 | p=0.005 |
| **Omnibus test for SES** | Pillai's Trace | 0.022 | F(3,63)=0.482 | p=0.696 |
| | Wilk's Lambda | 0.978 | F(3,63)=0.482 | p=0.696 |
| **Between Subjects Effects for Group** | sMRI | n/a | F(2,65)=3.937 | p=0.024 |
| | rsfcMRI | n/a | F(2,65)=2.927 | p=0.061 |
| | tfMRI | n/a | F(2,65)=5.895 | p=0.004 |
| **Post-hoc tests on estimated marginal means for Group** | **Item** | **Mean Difference** | **Standard Error** | **p-values** |
| | sMRI: HC - BP | 0.008 | 0.018 | p=0.638 |
| | sMRI: HC - SCZ | 0.054 | 0.020 | p=0.009 |
| | sMRI: BP - SCZ | 0.046 | 0.020 | p=0.026 |
| | rsfcMRI: HC - BP | -0.001 | 0.019 | p=0.958 |
| | rsfcMRI: HC - SCZ | 0.046 | 0.021 | p=0.035 |
| | rsfcMRI: BP - SCZ | 0.047 | 0.021 | p=0.032 |
| | tfMRI: HC - BP | 0.031 | 0.023 | p=0.171 |
| | tfMRI: HC - SCZ | 0.089 | 0.026 | p=0.001 |
| | tfMRI: BP - SCZ | 0.057 | 0.026 | p=0.030 |

**Table S10**: Cognitive control performance across groups with psychotic and non-psychotic BP split

| | Healthy Controls (HC) | Low Psychosis Bipolar (LPBP) | High Psychosis Bipolar (HPLP) | Schizophrenia (SZ) | Omnibus test statistic | p-values |
|---|---|---|---|---|---|---|
| **Cognitive Control Composite** | 0.44±0.51 (30) | 0.26±0.56 (15) | 0.01±0.70 (12) | -0.44±0.97 (23) | F(3,76) = 7.142 | Omnibus p<0.000<br>HC vs SZ p<0.000<br>HC vs LPBP p=0.847<br>HC vs HPBP p=0.281<br>SZ vs LPBP p=0.020<br>SZ vs HPBP p=0.292<br>LPBP vs HPBP p=0.792 |

**Caption:** BP participants were split into high and low psychotic symptom groups based on a median-split of the self-report psychosis score from the WERCAP.

Lerman-Sinkoff, D. B., J. Sui, S. Rachakonda, S. Kandala, V. D. Calhoun and D. M. Barch (2017). "Multimodal neural correlates of cognitive control in the Human Connectome Project." Neuroimage **163**: 41-54.
